# Chrom-pro: A User-Friendly Toolkit for De-novo Chromosome Assembly and Genomic Analysis

**DOI:** 10.1101/2024.03.02.583079

**Authors:** Wei Song, Tianrui Ye, Shaobo Liu, Dawei Shen, Yuhui Du, Yuening Yang, Yanming Lu, Hulin Jin, Yixin Huo, Weilan Piao, Hua Jin

## Abstract

Chromosome-level genome assembly is fundamental to current genomic and post-genomic research, however, the process remains complicated and challenging due to the lack of a standardized automatic workflow. The frequently-used method for high-quality genome assembly generally employs second-generation sequencing (SGS) low error reads, third-generation sequencing (TGS) long reads and Hi-C reads. In this study, we developed a multifunctional toolkit called Chrom-pro that integrated commonly-used algorithms for de novo chromosome-level genome assembly with above three data sets into a user-friendly, automatic workflow. Besides chromosome assembly, Chrom-pro also encompasses multiple functionalities for genome quality assessment, comparative genomic analysis, and structural variant detection, which offers substantial support for downstream research. To evaluate the performance of Chrom-pro software, we tested the software with publicly available sequencing data of mango, pufferfish, and plum, and the excellence was confirmed by achieving a BUSCO completeness score of over 95% as well as high collinearity with the reference genome. Furthermore, we applied Chrom-pro to investigating the impact of different internal algorithm options on the accuracy of chromosome assembly, providing guidance for advancing relevant research in the future. Overall, the development of Chrom-pro will significantly improve the efficiency and quality of chromosome assembly and contributing to the advancement of genomic research.

## Introduction

With the rapid development of high-throughput sequencing methods in the past two decades^1–4^, there has been a significant increase in genome assembly for various species. These assembled genomes have provided valuable insights into biological diversity, disease mechanisms, and evolution^5–9^. According to the lengths of assembled fragments, genomes can be broadly categorized into the contig/scaffold-level draft genome and the chromosome-level genome^10^. The high-quality chromosome-level genome is significantly important as it offers higher continuity and completeness^11, 12^, enabling comprehensive analysis of gene function and regulatory elements, precise interpretation of genetic information, as well as efficient genome engineering^13^.

There are two main approaches for genome assembly currently: reference-based assembly and de-novo assembly^14^. Reference-based assembly requires a reference genome from the same species or closely related species. In spite of advantages in efficiency and accuracy, reference-based assembly is hardly applied to the genomes unknown or from highly divergent species^15^. Common software for reference-based assembly includes RaGOO^14^, Ragout2^16^ and Metassembler^17^. The other approach, de-novo assembly, processes sequencing data directly, and is broadly applicable particularly for unknown genomes^18^. The long-read third-generation sequencing (TGS) can reduce the computation load of assembly and overcome the challenge posed by large repetitive sequences, making TGS competitive in resolving complex regions and structural variations of genomes, and suitable for de-novo assembly^19, 20^. Thus, Wtdbg2^21^, Flye^22^, and Canu^23^ have been developed specifically for de-novo assembly of long TGS reads into contigs. Furthermore, some new software HERA^24^ and GALA^25^ have recently emerged to process the assembled contigs together with TGS long reads to generate longer scaffolds, some of which can even reach the chromosomal level.

In order to achieve high-quality chromosome-level de-novo assembly, the most commonly-used approach integrates low-error second-generation sequencing (SGS), long-read TGS, and Hi-C sequencing^26–29^. Firstly, long TGS reads are assembled into contigs. Because of high error rates in TGS reads, assembled contigs are subsequently corrected using the more accurate SGS short reads^30^. Next, Hi-C sequencing is used to examine physical contact frequencies between different genomic regions, which provides information about the order and orientation of the assembled contigs to anchor them to chromosomes and complete chromosome-level genome assembly^31^. However, this widely-used strategy remains complicated because it requires manual step-by-step completion of most tasks without user-friendly software. This is time consuming and an obstacle for researchers with limited bioinformatics expertise^13^. In addition, the chromosomal assembly pipelines is inconsistent in different studies^27, 28^ and lack a well-tested, standardized workflow. Moreover, some critical steps such as removing haplotig, organelle DNA contamination and gap closing are prone to being overlooked ^32–34^. Given that only 0.17% of the eukaryotic species have undergone genome assembly^5^, there is a great need for a standardized and user-friendly automated tool for accurate chromosome-level genome assembly.

To address these issues, we developed Chrom-pro, the chromosome-level genome assembly software. Chrom-pro integrated commonly-used chromosome assembly software^27, 35, 36^ into a user-friendly and automatic workflow, achieving chromosome-level genome assembly. Furthermore, Chrom-pro provides several functional modules for downstream analysis, including genome quality assessment, result visualization, comparative genomics, and structural variation detection. We validated the performance of our software by reassembling genomes of three different species^37–39^ and comparing our results with the published reference genomes which were previously generated using various methods. In addition to these results, we also systematically evaluated the effects of different software on the accuracy of chromosome assembly results, providing guidance for choosing the optimal assembly software in future research.

## Methods

### Genome size estimation

Since the software for third-generation sequencing (TGS) reads assembly requires the genome size parameter, we provide two ways to enter genome size parameter. For the first way, we pre-process the raw reads to remove the adaptors and bases with low quality, by utilizing Fastp^40^ with standard parameters. Then, k-mer distribution is computed with jellyfish^41^, applying the parameters “-m 21 -C”, and the estimation of genome size is carried out using GenomeScope^42^. For the second way, the genome size can be measured by the flow cytometry method^43^, and then input to Chrom-pro using the “-genome size” parameter manually.

### Contig-level genome assembly

There are currently five available software options, Canu^23^, Flye^22^, Wtdbg2^21^, Hicanu^44^ and Raven^45^ for assembling TGS long reads from Nanopore and PacBio into contigs. Users can select one of the software for contig assembly by specifying the “-assemble_method” parameter. It is worth noting that Canu requires a large amount of memory and has relatively longer runtime. To address this issue, we extracted a 60X coverage of the sequencing data for contig assembly using Canu, while Flye and Wtdbg2 were executed with the entire sequencing data.

### Contig error correction

Due to the high error rate associated with TGS, it is necessary to correct the sequencing errors on the assembled contigs. Chrom-pro primarily integrated two error correction programs, namely Racon and Pilon. Initially, the assembled contigs undergo correction using “Racon v1.3.1”^46^ with TGS long reads. Subsequently, two rounds of polishing are performed using “Pilon v1.22”^47^ with SGS reads.

### Haplotig purging

The occurrence of haplotigs during the process of genome assembly may introduce redundancy that reduces the accuracy of the assembly. We employ the fully automatic tool purge_dups, which could be effectively integrated into the chromosome assembly pipeline. We use “purge_dups^48^ v1.1.2” with default parameters in Chrom-pro to identify and remove haplotypic duplication from primary genome assembly based on sequence similarity and read depth.

### Organelle genome removing

As the organelle genomes are usually sequenced together with nuclear genomes in eukaryotes, they need to be removed before assembling chromosomes. Chrom-pro first aligns the organelle genomes from NCBI database with all assembled contigs. If the alignment length is more than half of the assembled contig length and sequencing depth of the contig exceeds 20 times the average genome sequencing depth, the contig will be considered as organelle genomes and removed.

### Hi-C read mapping

HiC-Pro^49^ or HiCUP^50^ is utilized with default parameters to align Hi-C sequencing reads to de-novo assembled contigs and identify reliable ligation events. The restriction endonucleases used in Hi-C sequencing library construction need to be specified for running HiC-Pro or HiCUP. At present, the restriction endonucleases that can be accepted by Chrom-pro mainly include HindIII and MBOI.

### Chromosome anchoring

Chrom-pro offers two options, ALLHiC^51^ and LACHESIS^31^, for anchoring contigs into chromosomes. ALLHiC and LACHESIS cluster contigs into groups based on two principles: the frequency of intra-chromosomal interactions is significantly higher than that of inter-chromosomal interactions; the frequency of interactions on the same chromosome decreases with the increase in the interaction distance. The contigs in each group are further ordered and oriented to generate chromosomal level genomes.

### Gap closing

The initially-assembled chromosomes often possess Gaps, which are comprised of unknown bases and usually marked as ’N’s. These gaps possibly arise from factors like repetitive sequences, and their presence can hinder full understanding of the genome. Chrom-pro first runs abyss sealer^52^ software with SGS reads, and then runs TGS-GapCloser^53^ software with TGS reads to recover these unknown bases.

### Chromosome visualization

Chrom-pro first calculates the GC content of each chromosome by seqkit^54^. Then, BWA^55^, minimap2^56^, and HiCUP are used to align SGS, TGS and Hi-C reads to chromosomes respectively, obtaining the sequencing depth of different sequencing methods. JCVI^57^ software is used to compute the collinearity blocks between chromosomes. Finally, circos^58^ is used to visualize chromosome length, GC content, sequencing depth, and collinearity blocks. The command for chromosome visualization is shown below:

docker run -v /var/run/docker.sock:/var/run/docker.sock -v $(pwd):/data -w /data songweidocker/genomic_analysis:v1 python /usr/bin/circos_plot.py -TGS TGS.fastq -NGS_1 NGS_1.fastq -NGS_2 NGS_2.fastq -hic_1 HiC_1.fastq -hic_2 HiC_2.fastq -hic_Enzyme HINDIII -threads 8 -genome assembled.fasta

### Comparative genomics

JCVI software is used to carry out the collinearity analysis between the assembled chromosomes and reference genomes. Due to the limited capability of JCVI in visualizing chromosomal co-linearity and the absence of quantitative analysis for inter-chromosomal rearrangements, we developed a custom script that allows to quantitatively assess the inversion and translocation events between different chromosomes based on the output of JCVI. The command for comparative genomics is shown below:

docker run -v /var/run/docker.sock:/var/run/docker.sock -v $(pwd):/data -w /data songweidocker/genomic_analysis:v1 python /usr/bin/genome_comepare.py assembled_genome.fasta reference_genome.fasta

### Genome structural variation detection

Prior to detecting structural variations between two versions of chromosomes using SyRI, it is essential to ensure that the homologous chromosomes in the two genomes display the same strand sequences. The tools like seqkit can achieve maintaining strand consistency between a chromosome pair. Chrom-pro utilizes SyRI^59^ software to detect various structural variations, such as SNPs, Indels, duplications, and translocations, between different versions of chromosomes. The identified collinear regions and structural variations between different versions can be visualized using Plotsr^60^ software. The command for structural variation detection is shown below:

docker run -v /var/run/docker.sock:/var/run/docker.sock -v $(pwd):/data -w /data songweidocker/genomic_analysis:v1 python /usr/bin/syri_plotsr.py assembled_genome.fasta reference_genome.fasta

### The quality assessment of chromosome assembly

After completing chromosome assembly, Chrom-pro will utilize Quast^61^ software to generate statistics on the longest contig length, contig N50, and the number of gaps; then BUSCO^62^ software to evaluate the completeness of the assembled chromosomes; Eukaryota_odb10 as the default database to assess the integrity. The command for genome quality assessment is shown below:

docker run -v /var/run/docker.sock:/var/run/docker.sock -v $(pwd):/data -w /data songweidocker/genomic_analysis:v1 python /usr/bin/busco_calculate_plot.py assembled_genome.fasta

### Masking repetitive sequences on genome

Repetitive sequences are identified using the combination of homology-based and de-novo methods. Firstly, potential LTR retrotransposons are identified using LTRharvest^63^ and LTR_Finder^64^. The results from these programs are then integrated and refined with LTR_retriever to provide a high-confident set of LTR retrotransposons. In parallel, RepeatModeler^65^ is employed to identify the spectrum of repeats across the genome sequences. Lastly, both output from LTR_retriever and RepeatModeler comprise a library of repetitive sequences. According to this library, all repetitive regions across the genome are masked using RepeatMasker^66^ software.

### Availability of data and materials

The Docker image of Chrom-pro is deposited at Docker hub https://hub.docker.com/repository/docker/songweidocker/genomic_analysis/. All tests were performed on an Ubuntu Linux release 18.04.3 server with two Intel Xeon processors (3 2 cores each, 64 threads total) and 512 GB of RAM. The *Prunus salicina* (plu m) sequencing dataset is available at https://www.ncbi.nlm.nih.gov/bioproject/PRJNA574159. The *Takifugu bimaculatus* (pufferfish) sequencing data-set is availab le at https://www.ncbi.nlm.nih.gov/bioproject/PRJNA508537. The *Mangifera indic a* (mango) sequencing dat-aset isavailable at https://www.ncbi.nlm.nih.gov/bioproject/PRJNA487154/.

## Results

### 1. The overview of multifunctional toolkit Chrom-pro

#### 1.1 Chromosome-level genome assembly with Chrom-pro

The primary function of Chrom-pro is chromosome-level genome assembly, which requires three data sets as input: SGS (also known as next-generation sequencing, NGS) reads, TGS reads, and Hi-C reads (Fig. 1A). Prior to assembly, Chrom-pro employs internal programs, Jellyfish^41^ and GenomeScope^42^, to estimate genome size with the NGS reads and passes the results to downstream contig assembly software. Alternatively, if the genome size is already known from other methods like flow cytometry, it can be directly specified using the “-genome size” parameter. To assemble long TGS reads into contigs in the next step, Chrom-pro offers three possible options, Wtdbg2^21^, Flye^22^, and Canu^23^. Due to the high error rate in TGS reads, Chrom-pro corrects the assembled contigs using both TGS and NGS reads with Racon^46^ and Pilon^47^ software respectively. After error correction, haplotigs are removed from the contigs with the Purge_dups^48^ in Chrom-pro. To further remove the organelle genome sequences, the organelle genomes from NCBI database are aligned with all contigs. Then, the organelle-genome-derived contigs are detected and removed according to the alignment result and sequencing depths of the contigs, as the copy number of organelle genome is practically much higher than that of nuclear genome in a cell. Hi-C reads are then aligned to the filtered contigs using either HiC-Pro^49^ or HiCUP^50^. The contigs are clustered, sorted, and oriented with LACHESIS^31^ or ALLHiC^51^, based on the frequency of Hi-C interactions between contigs to obtain chromosomal level genomes. Finally, Chrom-pro performs gap-closing on the assembled chromosomes by successively running Abyss-sealer inputted with SGS reads and TGS-GapCloser^53^ inputted with TGS reads to refine the chromosome assembly. The entire assembly process can be executed by entering a single command in Chrom-pro (Fig. 1B), which will significantly simplify the process of chromosome assembly.

**Figure 1.**
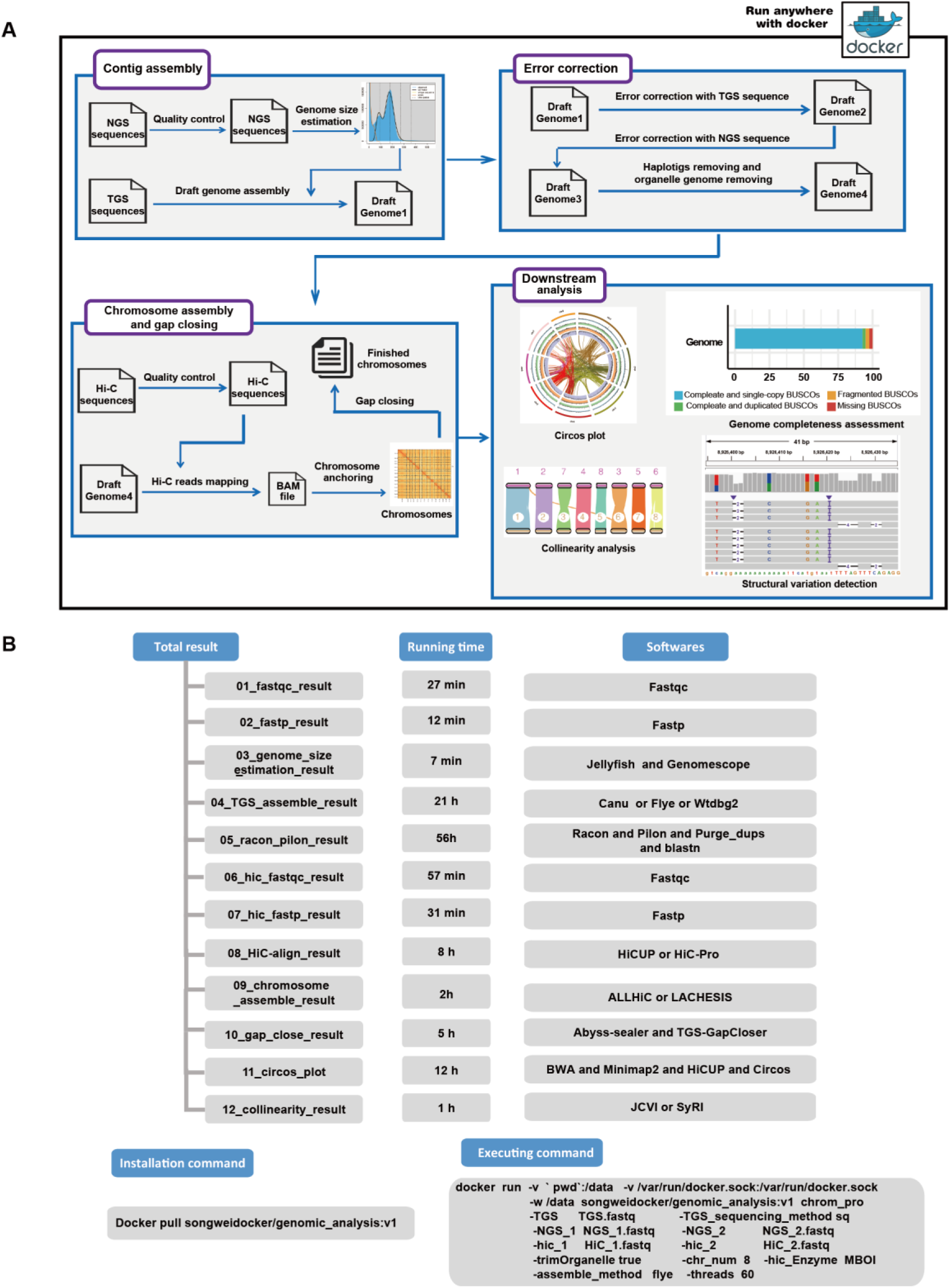
Workflow of the Chrom-pro software package. **(A)** Schematic overview of the included function and overall running procedure in Chrom-pro. **(B)** The required software, running time and obtained result in each step of Chrom-pro. The running time is estimated based on the results of mango genome assembly.

#### 1.2 Assessment and visualization of assembled chromosomes with Chrom-pro

Chromosome visualization is achieved through the Circos software ^58^, which generates a circular map displaying chromosome length, GC content, and sequencing depth of SGS, TGS and Hi-C reads as well as the information about collinearity between chromosomes (Fig. 2A-2C). To assess the quality of chromosome assembly result, Chrom-pro incorporated two powerful tools: Quast^61^ and BUSCO (Benchmarking Universal Single-Copy Ortholog)^62^. Quast can compute various metrics including contig number, contig N50, gap number, and the length of the longest contig (Table 1). On the other hand, BUSCO can evaluate the completeness and accuracy of assembled chromosomes using the Eukaryota_odb10 database by default (Fig. 2J-2L). Users can also run BUSCO independently and customize the database using the “-db” parameter. The better assembly is typically characterized by longer contig N50, fewer contig numbers, and higher BUSCO scores. These functions produce a comprehensive assessment of the quality of chromosome assembly and support researchers to make appropriate decisions on downstream analysis.

**Figure 2.**
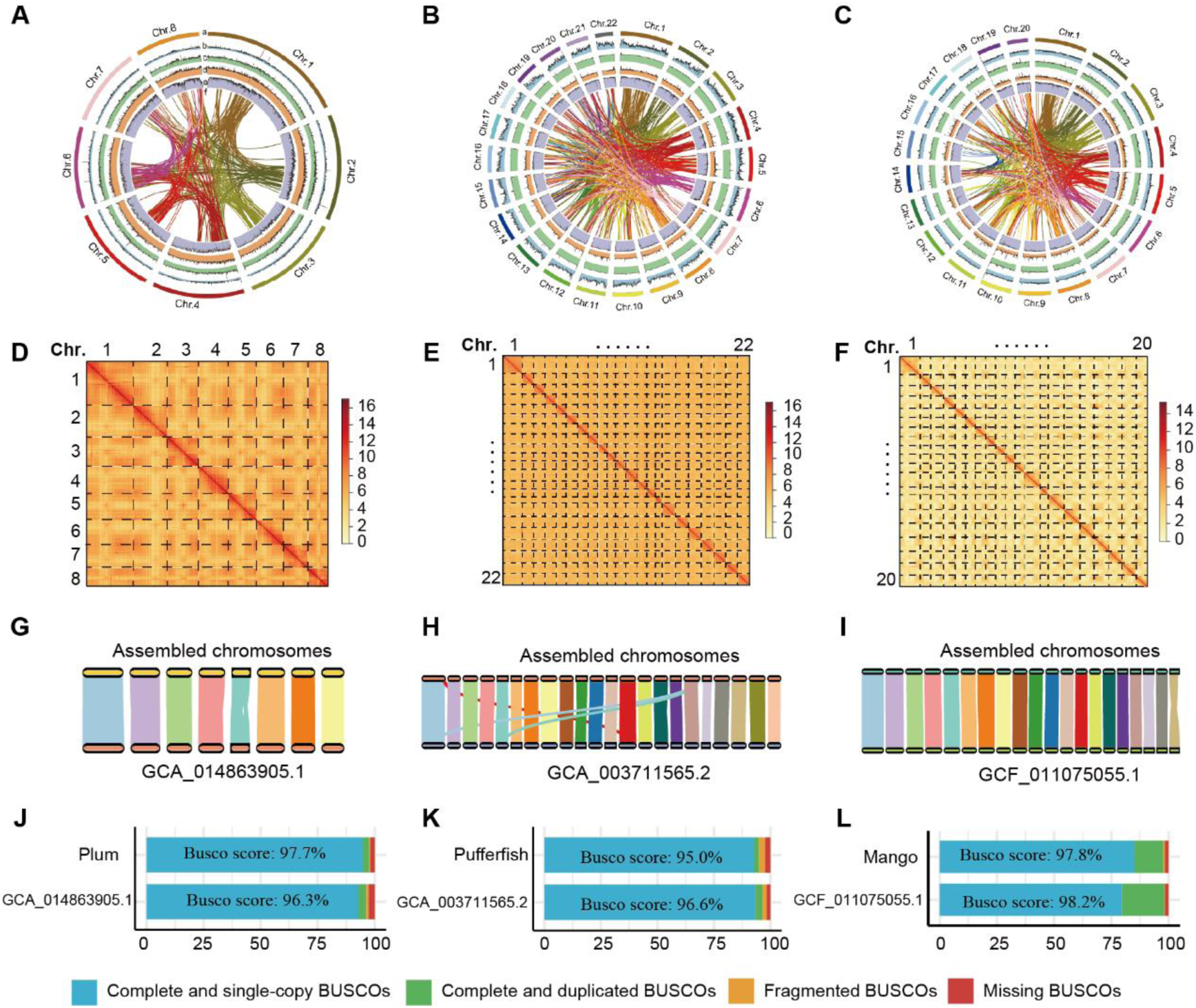
Chromosome assembly results of plum, pufferfish, and mango. **(A-C)** Circular chromosome maps for plum (A), pufferfish (B), and mango (C). From outer to inner rings: chromosome, GC content, SGS sequencing depth, TGS sequencing depth, Hi-C sequencing depth, and collinearity block between chromosomes. **(D-F)** Hi-C contact frequency heat map of assembled chromosomes for plum (D), pufferfish (E), and mango (F). **(G-I)** The collinearity analysis results between the assembled chromosomes (top) and the corresponding NCBI reference genomes (bottom) from plum (G), puffer fish (H), and mango (I). **(J-L)** BUSCO completeness evaluation for the assembled (top) and reference (bottom) genomes of plum (J), pufferfish (K), and mango (L).

**Table 1.** Comparison of chromosome assemblies conducted with different software combinations for Plum, Pufferfish, and Mango. The contig assembly was performed with Canu, Wtdbg2, or Flye; Hi-C reads mapping was carried out with HiC-Pro or HiCUP; chromosome anchoring was done with ALLHiC or LACHESIS. The primary contig number reflects the total contig number initially generated by contig-assembly software; the purged contig number represents the contig number after eliminating haplotig duplication from primary contigs. The completeness of assembled genome was assessed using BUSCO.

| Species and assembly method | Primary contig number | Primary contig N50 | Purged contig number | Purged contig N50 | Assembly size | % of assembly in pseudomolecules | BUSCO score |
| --- | --- | --- | --- | --- | --- | --- | --- |
| Plum_Canu_HiC-Pro_LACHESIS | 7397 | 99.3Kb | 2807 | 217.0Kb | 285.3Mb | 98.65 | 97.40% |
| Plum_Canu_HiC-Pro_ALLHiC | 7397 | 99.3Kb | 2807 | 217.0Kb | 282.93Mb | 99.32 | 97.60% |
| Plum_Canu_HiCUP_LACHESIS | 7397 | 99.3Kb | 2807 | 217.0Kb | 285.21Mb | 95.67 | 97.50% |
| Plum_Canu_HiCUP_ALLHiC | 7397 | 99.3Kb | 2807 | 217.0Kb | 282.93Mb | 99.42 | 97.20% |
| Plum_Flye_HiC-Pro_LACHESIS | 4097 | 200.0Kb | 2925 | 222.2Kb | 274.93Mb | 99.25 | 97.70% |
| Plum_Flye_HiC-Pro_ALLHiC | 4097 | 200.0Kb | 2925 | 222.2Kb | 272.74Mb | 99.92 | 97.70% |
| Plum_Flye_HiCUP_LACHESIS | 4097 | 200.0Kb | 2925 | 222.2Kb | 274.91Mb | 98.45 | 97.90% |
| Plum_Flye_HiCUP_ALLHiC | 4097 | 200.0Kb | 2925 | 222.2Kb | 272.73Mb | 99.87 | 97.70% |
| Plum_Wtdbg2_HiC-Pro_LACHESIS | 14462 | 38.9Kb | 9202 | 50.3Kb | 341.88Mb | 83.04 | 81.90% |
| Plum_Wtdbg2_HiC-Pro_ALLHiC | 14462 | 38.9Kb | 9202 | 50.3Kb | 335.45Mb | 95.26 | 82.40% |
| Plum_Wtdbg2_HiCUP_LACHESIS | 14462 | 38.9Kb | 9202 | 50.3Kb | 341.45Mb | 79.1 | 81.90% |
| Plum_Wtdbg2_HiCUP_ALLHiC | 14462 | 38.9Kb | 9202 | 50.3Kb | 335.42Mb | 93 | 82.10% |
| Pufferfish_Canu_HiC-Pro_LACHESIS | 4957 | 329.2Kb | 1970 | 517.1Kb | 358.15Mb | 99.81 | 94.80% |
| Pufferfish_Canu_HiC-Pro_ALLHiC | 4957 | 329.2Kb | 1970 | 517.1Kb | 356.44Mb | 99.99 | 95.00% |
| Pufferfish_Canu_HiCUP_LACHESIS | 4957 | 329.2Kb | 1970 | 517.1Kb | 358.14Mb | 99.81 | 94.80% |
| Pufferfish_Canu_HiCUP_ALLHiC | 4957 | 329.2Kb | 1970 | 517.1Kb | 356.44Mb | 99.99 | 94.80% |
| Pufferfish_Flye_HiC-Pro_LACHESIS | 1485 | 2833.3Kb | 614 | 2917.9Kb | 359.47Mb | 99.26 | 96.20% |
| Pufferfish_Flye_HiC-Pro_ALLHiC | 1485 | 2833.3Kb | 614 | 2917.9Kb | 359.08Mb | 99.44 | 96.30% |
| Pufferfish_Flye_HiCUP_LACHESIS | 1485 | 2833.3Kb | 614 | 2917.9Kb | 359.47Mb | 99.6 | 96.40% |
| Pufferfish_Flye_HiCUP_ALLHiC | 1485 | 2833.3Kb | 614 | 2917.9Kb | 359.08Mb | 99.85 | 96.50% |
| Mango_Canu_HiC-Pro_LACHESIS | 2793 | 499.2Kb | 596 | 1112.8Kb | 358.24Mb | 99.72 | 97.80% |
| Mango_Canu_HiC-Pro_ALLHiC | 2793 | 499.2Kb | 596 | 1112.8Kb | 357.73Mb | 99.81 | 97.80% |
| Mango_Canu_HiCUP_LACHESIS | 2793 | 499.2Kb | 596 | 1112.8Kb | 358.23Mb | 99.65 | 97.80% |
| Mango_Canu_HiCUP_ALLHiC | 2793 | 499.2Kb | 596 | 1112.8Kb | 357.73Mb | 99.95 | 97.70% |
| Mango_Flye_HiC-Pro_LACHESIS | 2544 | 792.0Kb | 1433 | 919.3Kb | 367.05Mb | 99.67 | 97.80% |
| Mango_Flye_HiC-Pro_ALLHiC | 2544 | 792.0Kb | 1433 | 919.3Kb | 365.98Mb | 99.99 | 97.80% |
| Mango_Flye_HiCUP_LACHESIS | 2544 | 792.0Kb | 1433 | 919.3Kb | 367.06Mb | 99.74 | 97.70% |
| Mango_Flye_HiCUP_ALLHiC | 2544 | 792.0Kb | 1433 | 919.3Kb | 365.98Mb | 99.99 | 97.80% |

#### 1.3 Comparative genomics analysis with Chrom-pro

Chrom-pro is a useful tool that not only facilitates the assembly of chromosomes, but also offers comparative genomics functionality to reveal collinearity and structural variations between assembled chromosomes and other reference genomes. Chrom-pro employs the widely-used JCVI ^57^ tool for detecting chromosomal collinearity (Fig. 2G-2I). Single nucleotide polymorphism (SNP), insertion and/or deletion (Indel) and other structural variations can be detected with the SyRI^59^ and visualized using the plotsr ^60^ (Fig. 3A). With these comprehensive features, Chrom-pro will be a valuable tool for researchers seeking to explore the complex and dynamic genomic landscape.

**Figure 3.**
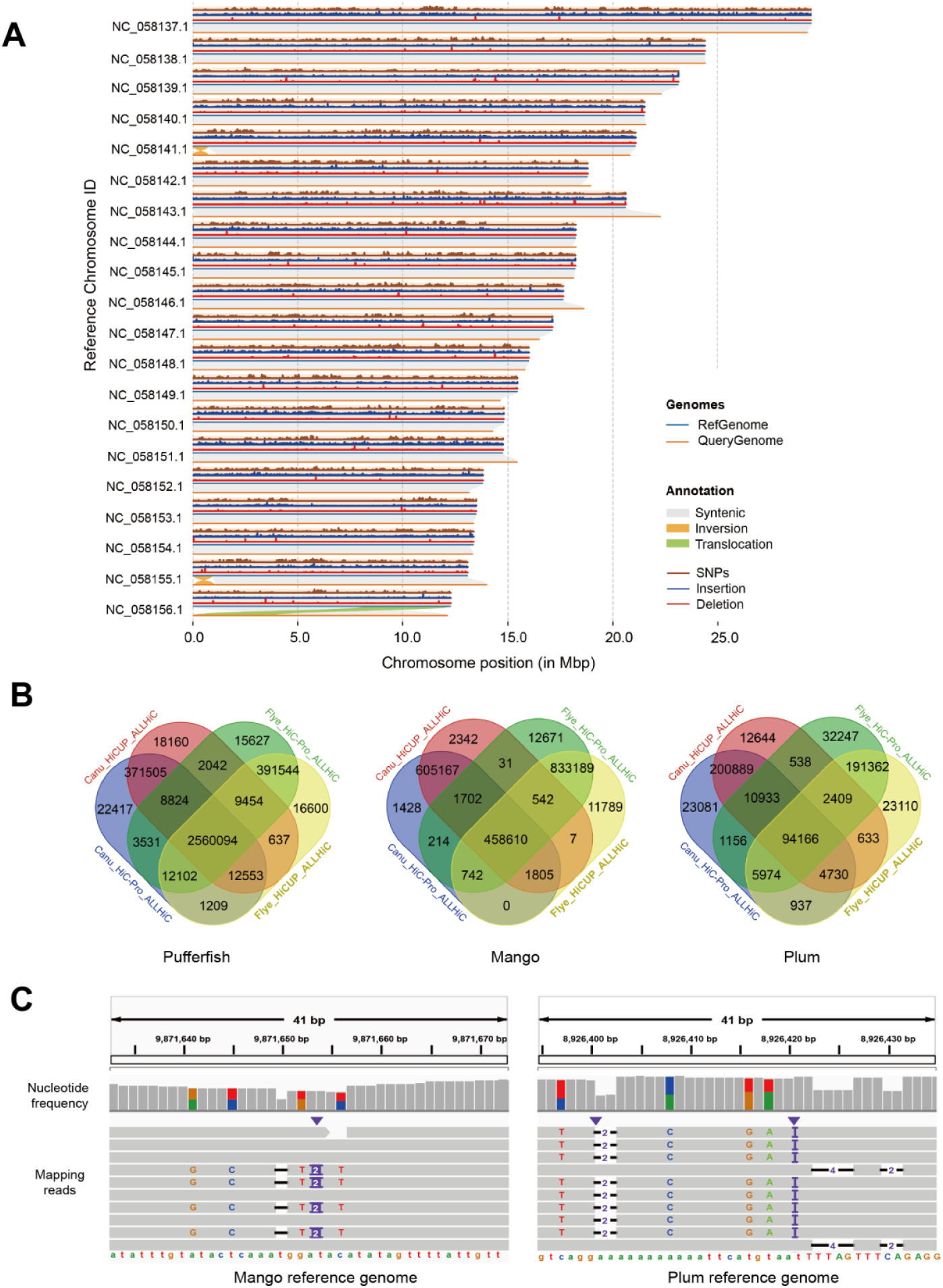
Comparative genomic analysis of Chrom-pro reveals SNP, Indel, and SV between assembled and reference genomes. **(A)** Collinear regions and variations between assembled chromosomes and reference genomes of mango. **(B)** The SNPs between chromosomes assembled by distinct combination of software. **(C)** Validation of SNPs and Indels through SGS reads mapping. SNP: single nucleotide polymorphism, Indel: insertion and deletion, SV: structural variation.

### 2. The performance of Chrom-pro in the chromosome assembly of plum, puffefish, and mango

To evaluate the accuracy of Chrom-pro, we utilized publicly available sequencing data of *Prunus salicina* (plum)^37^, *Takifugu bimaculatus* (pufferfish)^38^ and *Mangifera indica* (mango)^39^. These species have multiple versions of chromosome-level genomes in the NCBI database, enabling accurate comparison with Chrom-pro-assembled chromosomes. Plum, Pufferfish, and mango have 8, 22, and 20 chromosomes respectively. Table S1 listed the sequencing methods, data sizes, and chromosome assembly methods used in the published research and Table S2 listed the assembly options used in Chrom-pro.

After chromosome assembly, we evaluated the accuracy of Chrom-pro output in four ways. Firstly, we generated heat map of Hi-C interaction between different positions in the assembled chromosomes using ALLHiC in Chrom-pro (Fig. 2D-F). The heat maps show that the Hi-C interaction frequencies between adjacent positions are higher than those between distant positions, with minimal uneven interaction frequencies within and between chromosomes (Fig. 2D-F). Secondly, the Chrom-pro-assembled chromosomes showed high collinearity with their corresponding NCBI reference genomes (Fig. 2G-I). Despite some displacement events appearing near chromosome ends in pufferfish, these likely arise from repetitive sequences in telomeric regions. Thirdly, the chromosome completeness was assessed with BUSCO using the Eukaryota_odb10 database (Fig. 2J-L). Over 95% of BUSCO genes were identified in our assembled genomes, demonstrating the accuracy and reliable of the assembled chromosome for downstream analysis. Fourthly, more than 98% of original TGS and SGS reads could be mapped back to the assembled chromosomes. Altogether, these evaluations convincingly demonstrated Chrom-pro as a reliable tool for high-quality chromosome-level genome assembly.

Next, we employed SyRI within Chrom-pro to detect single nucleotide polymorphism (SNP), insertion and deletion (Indel) and structural variation (SV) between the assembled and reference genomes. Plotsr in Chrom-pro was then utilized to visualize the events of synthetic, inversion, and translocation, as well as the frequency of SNP, insertion, and deletion (Fig. 3A). Subsequently, we compared the SNP between the reference genomes and the chromosomes assembled by four distinct combinations of software, finding that 98%, 99%, and 85% of SNP in pufferfish, mango, and plum could be reproducibly detected at least three times out of four (Fig. 3B). To further validate the accuracy of these identified SNPs, we aligned the SGS reads to the reference genomes. This alignment confirmed that over 80% of SNPs in mango and pufferfish, and 60% of SNPs in plum could be verified by the SGS reads. As shown in Figure 3C, multiple reads support the variation at each position, suggesting that the SNPs and SVs detected by Chrom-pro are accurate.

### 3. The impact of different software on the accuracy of chromosome assembly

Accurate chromosome assembly is essential for post-genomic research, while studies focusing on the impact of different software on the accuracy of chromosome assembly are still limited. To explore the impact of various software tools used in different stages of chromosome assembly, we utilized Chrom-pro in conjunction with multiple software combinations to assess their impact on the accuracy of chromosome assembly.

#### 3.1 Comparison of Wtdbg2, Flye, and Canu for contig assembly

Since contig assembly forms the foundation of chromosome assembly, we compared the impact of three widely-used contig assembly software —Wtdbg2, Flye, and Canu— on the accuracy of chromosome assembly in plum, pufferfish, and mango. Our results revealed that Flye and Canu were successful in assembling chromosomes of all three species, while Wtdbg2 was only successful in plum but not in pufferfish and mango (Table 1). Furthermore, 99% of contigs assembled by Flye and Canu were finally anchored to chromosomes. In contrast, Wtdbg2 showed a slightly lower anchoring rate of 88%. According to genome completeness assessed by BUSCO, both Flye and Canu achieved more than 95% of assembly completeness, whereas Wtdbg2 only reached 82% (Table 1). According to the collinearity analysis result, it was obvious that the assembly by Canu and Flye demonstrated higher consistency with the reference genomes than that by wtdbg2 (Figure 5).

**Figure 4.**
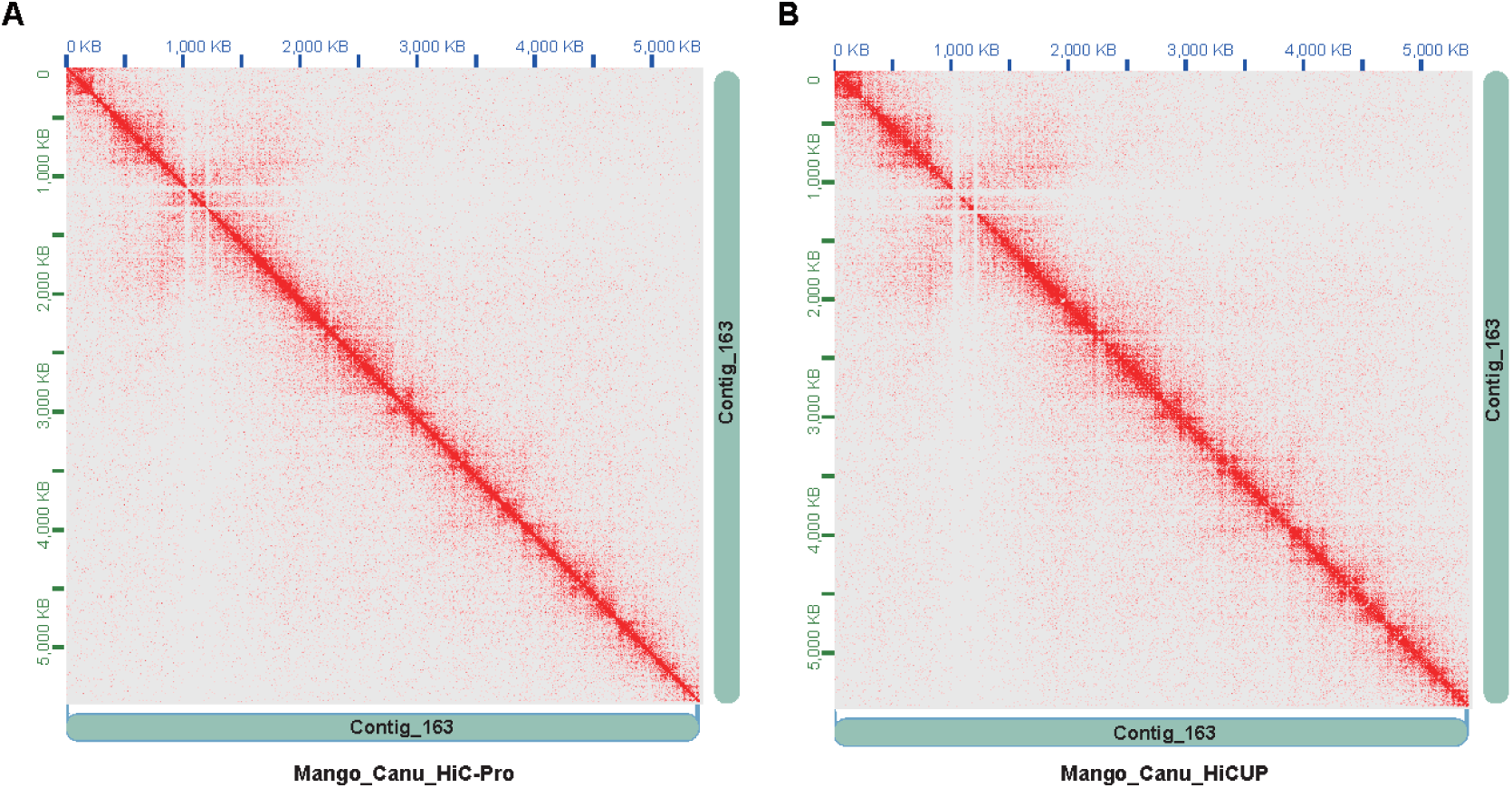
Hi-C contact frequency maps generated by HiC-Pro (A) and HiCUP (B).

**Figure 5.**
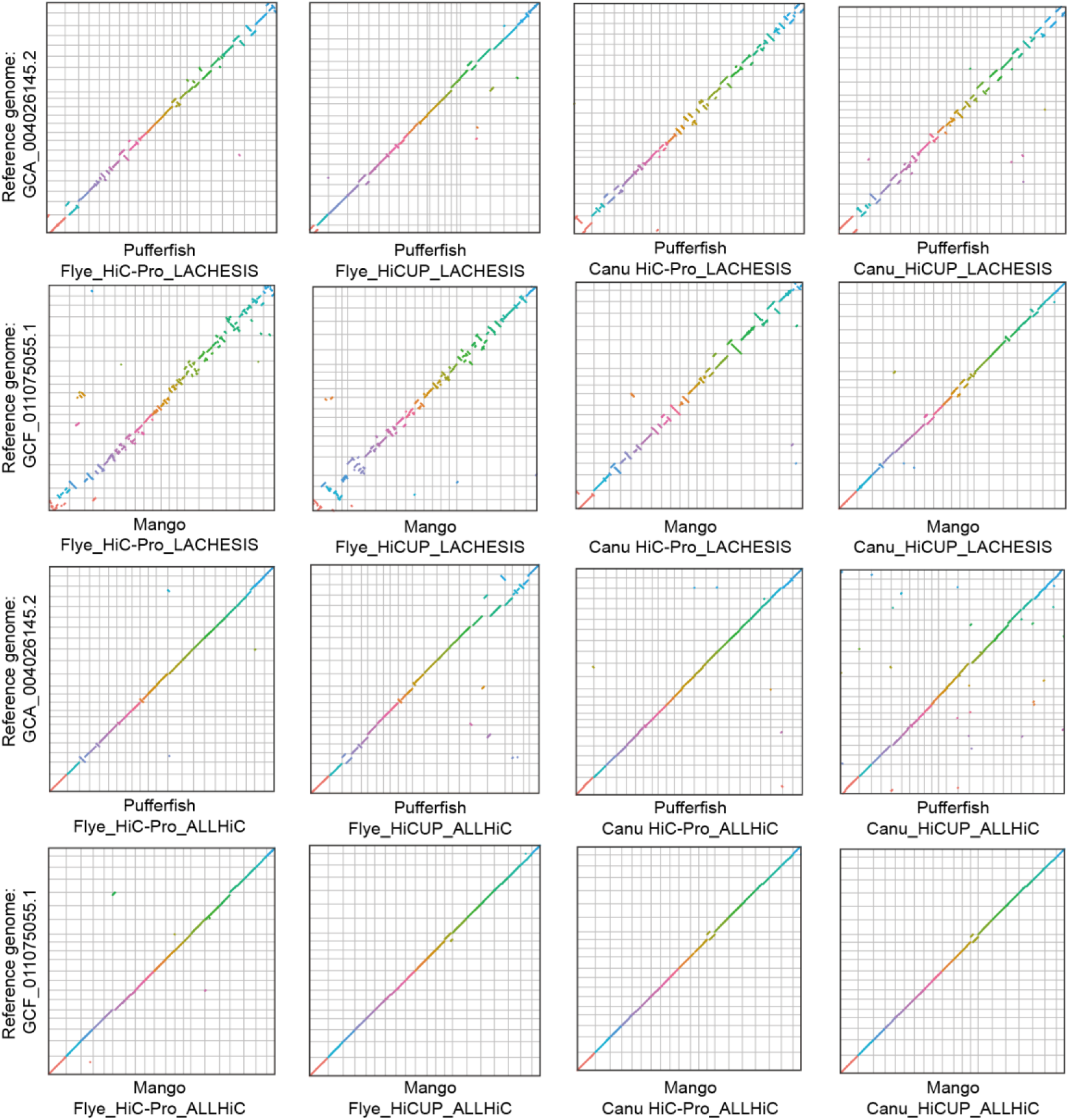
Collinearity analysis between assembled chromosomes and corresponding reference genomes from the NCBI genome database. The vertical and horizontal axes represent the assembled chromosomes and their respective reference genomes. Distinct chromosomes are indicated by different colors.

#### 3.2 Comparison of HiC-Pro and HiCUP for Hi-C read mapping

Hi-C sequencing has become necessary for genome assembly, primarily due to its capability to align and anchor disordered contig sequences to create chromosome-level genome. This effectiveness stems from the characteristic of Hi-C data, where the interaction strength between chromatin segments decreases as the distance between them increases. Consequently, segments that are closer together tend to exhibit a higher frequency of interactions. HiC-Pro and HiCUP are two widely used software for processing Hi-C data. HiC-Pro searches for restriction enzyme sites in Hi-C reads that have either failed to align to the assembled contigs (unmapped) or aligned to multiple locations (multi-mapped). Then, HiC-Pro splits these reads at the ligated positions (restriction enzyme sites) into two pieces and remaps these pieces to the assembled contigs separately. On the other hand, HiCUP truncates all raw Hi-C reads at the ligated positions if present, and then aligns the truncated pieces to the assembled contigs using bowtie2^67^.

According to the comparison of mapping results produced by HiC-Pro and HiCUP, wefound that the alignment percentages of valid pairs and uniquely valid pairs (non-redundant, true ligation products) from HiC-Pro were close to those from HiCUP (Table 2). Additionally, the Hi-C contact frequency maps from the two programs showed similar interaction patterns (Fig. 4A & 4B), indicating comparable effectiveness of HiC-Pro and HiCUP in processing Hi-C data. Furthermore, we also observed no significant advantage of either software in terms of chromosome collinearity analysis (Fig. 5 and Fig. S1). Therefore, these results indicated that both HiC-Pro and HiCUP can properly carry out Hi-C read mapping and chromosome assembly with similar effectiveness.

**Table 2.** Comparison of valid and uniquely valid pairs generated by HiC-Pro and HiCUP. Valid pairs refer to paired reads from Hi-C sequencing that are properly mapped and have a correct ligation site, while uniquely valid pairs refer to a subset of valid pairs where both reads map to unique locations in the genome.

|  | HiC-Pro |  | HiCUP |  |
| --- | --- | --- | --- | --- |
|  | Valid pairs | Uniquely valid pairs | Valid pairs | Uniquely valid pairs |
| Mango-Canu | 36% | 32% | 33.30% | 30% |
| Mango-Flye | 37% | 33% | 33.50% | 30% |
| Pufferfish_Canu | 65% | 32% | 71.90% | 35% |
| Pufferfish_Flye | 65% | 32% | 72.30% | 35% |
| Plum_Canu | 94% | 80% | 92% | 78% |
| Plum_Flye | 94% | 80% | 92.20% | 78% |
| Plum_Wtdbg2 | 93% | 79% | 90.30% | 76% |

#### 3.3 Comparison of LACHESIS and ALLHiC for anchoring contigs to chromosome based on Hi-C sequencing data

Currently, LACHESIS and ALLHiC are the two major software used to anchor contigs to chromosome based on the interaction frequency matrix of Hi-C sequences. Both LACHESIS and ALLHiC utilize hierarchical clustering method to cluster contigs into user-defined chromosome groups^68^. Subsequently, LACHESIS uses the minimum spanning tree criterion to order and locate the clustered contigs, while ALLHiC uses a genetic algorithm to optimize the ordering and positioning of clustered contigs^51^. Collinearity analysis between the reference genome and the chromosomes anchored by ALLHiC or LACHESIS revealed that, under the same condition, the reference genome had a significantly higher collinearity with the chromosomes anchored by ALLHiC than LACHESIS (Fig. 5 and Fig. S1). Further examination revealed that although both tools can accurately cluster contigs into a specific number of chromosomes, ALLHiC was more accurate in ordering and orienting these clustered contigs compared to LACHESIS (Fig. 5 and Fig. S1). Moreover, statistical analysis of the structural discrepancies between the reference genome and assembled chromosomes uncovered that the number of inversion and translocation events was much less in the chromosomes anchored by ALLHiC compared to those by LACHESIS (Table 3).

**Table 3.**
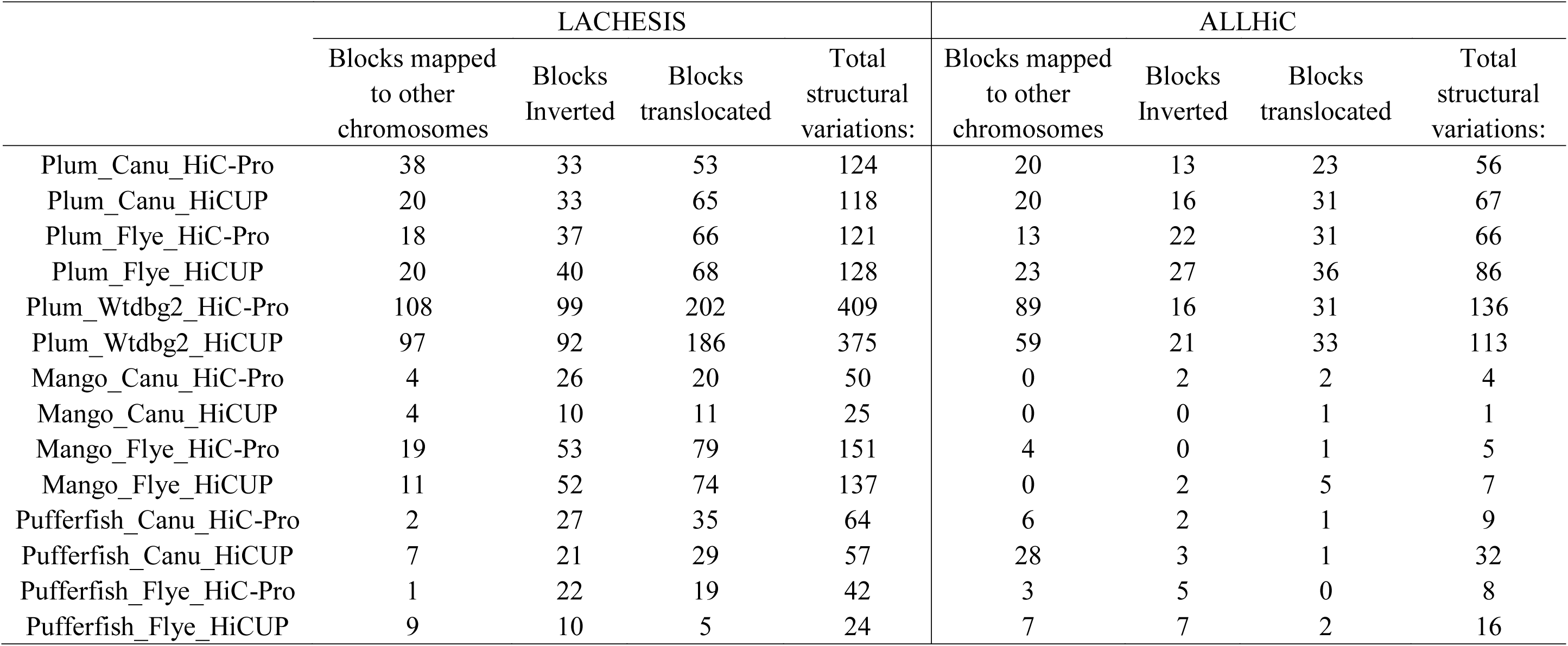
Statistical analysis of structural discrepancies between the reference genome and chromosomes assembled using various software combinations. Canu, Flye, and Wtdbg2 are applied for contig assembly; HiC-Pro and HiCUP are used for aligning Hi-C sequences to contigs; LACHESIS and ALLHiC are utilized for anchoring contigs into chromosomes.

#### 3.4 The influence of contig continuity and repetitive sequences on the accuracy of LACHESIS in chromosome anchoring

LACHESIS is one of the most widely used software developed for chromosome anchoring of contigs. Given its wide application, identifying factors that influence the accuracy of LACHESIS is critical, as this step directly impacts the quality of genome assemblies. By comparing the collinearity between reference genome and the chromosomes anchored by LACHESIS, we found that when the N50 of the contigs was larger, the chromosomes anchored by LACHESIS had higher consistency with the reference genome (Fig. 6), as suggested by previous research^51^. On the other hand, we also found that LACHESIS tended to incorrectly anchor contigs in genomic regions with high repetitive sequence content (Fig. 6), implying that high repetitive sequence content may also affect the accuracy of LACHESIS in anchoring contigs.

**Figure 6.**
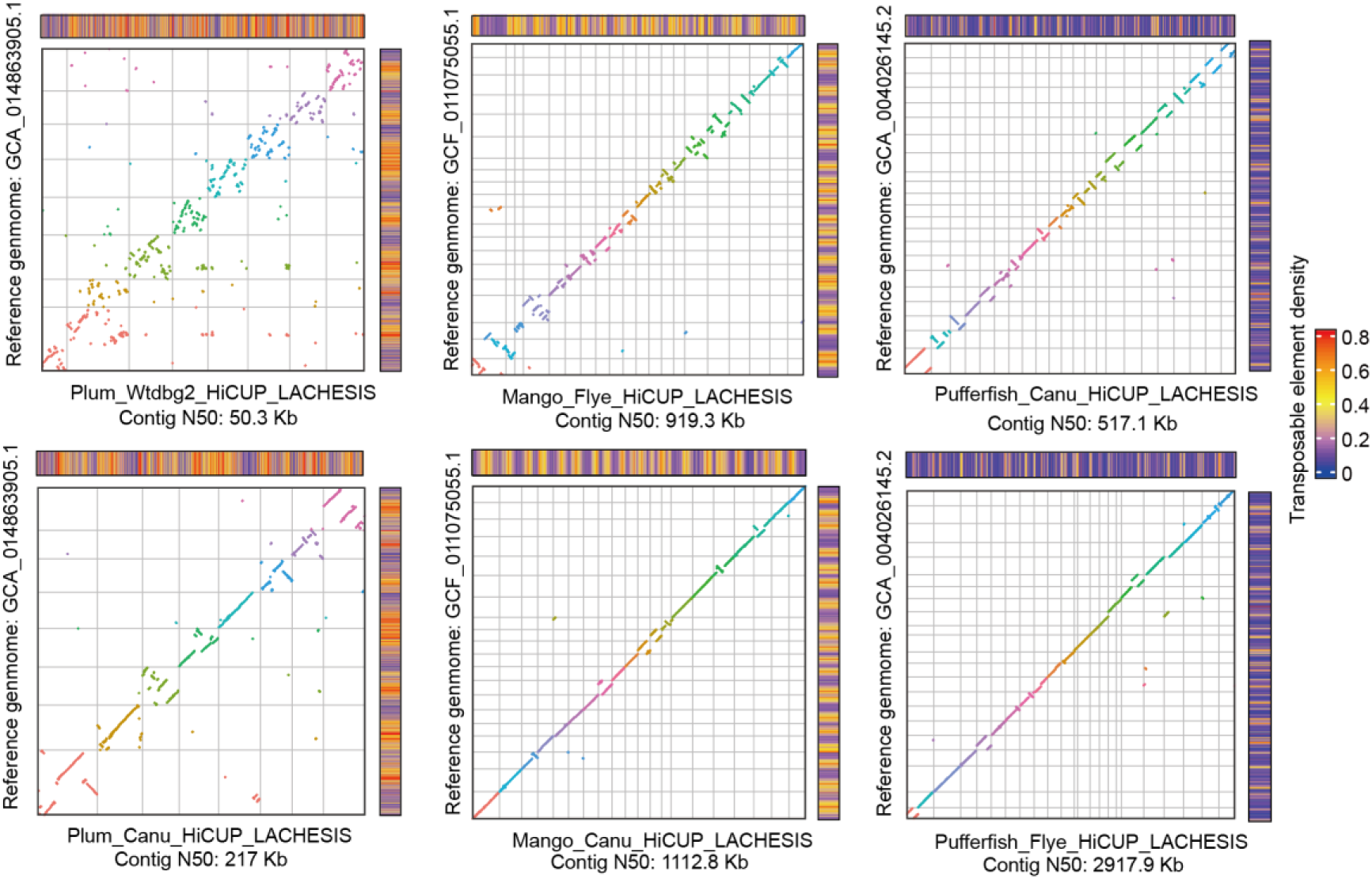
Influence of repetitive sequences and contig N50 on the anchoring accuracy of LACHESIS software. The vertical and horizontal axes represent the LACHESIS-anchored-chromosomes and the corresponding reference genomes. The heatmap represents the density of repetitive sequences on both the assembled chromosomes and reference genomes.

## Discussion

Although the strategy combining TGS, SGS and Hi-C to assemble a chromosome-scale genome has been gradually accepted, this widely used strategy remains complicated, requiring manual step-by-step completion of most tasks without user-friendly automated software. This is not only time-consuming but also prone to errors, particularly when dealing with large amounts of data. Given that the majority of species still lack chromosome-level genomes^5^ available for use, there is an urgent need for user-friendly and automatic software to accurately assemble chromosome-level genomes.

To address these issues, we developed Chrom-pro, an automated software specifically designed for chromosome-level genome assembly. Chrom-pro holds the following merits: (I) Chrom-pro is a user-friendly tool, easy to install and use, also capable of automatically assembling high-quality chromosome-level genomes. (II) In addition to genome assembly function, Chrom-pro also possesses the incorporated functionalities relevant to genome quality assessment, comparative genomic analysis, and structural variation detection. (III) The algorithm hired at each step of Chrom-pro is not static, offering various software options to ensure flexibility. Varying available software combinations not only enhance the success rate in assembly but also allow the cross-verification of assembled chromosomes. (IV) By implementing the pipeline within a Docker container, Chrom-pro has achieved a simplified and portable process for chromosome assembly and genomics analysis, facilitating distribution and execution across diverse computing platforms. Overall, the development of Chrom-pro fills a technical gap in this field and will significantly improve the efficiency and quality of chromosome assembly.

To validate the performance of Chrom-pro in chromosome assembly, we utilized Chrom-pro to assemble genomes with sequencing data from pufferfish, mango, and plum, and compared them with publicly available reference genomes. The genome completeness evaluated by BUSCO software indicates that Chrom-pro-assembled chromosomes achieve completeness over 95%, which closely matches the completeness of the reference genomes. Chromosomal collinearity analysis also demonstrates a high degree of collinearity between the chromosomes assembled by Chrom-pro and the reference genomes. These results reveled the high completeness and accuracy of chromosomes assembled by Chrom-pro, confirming its accuracy and reliability for chromosome assembly.

Accurate chromosome assembly is fundamental for post-genomic research, while few studies have focused on the impact of different software on the accuracy of final assembly outcomes^69^. In this study, we executed Chrom-pro with diverse software combination options for chromosome assembly. This process yielded 12 plum genomes, 8 pufferfish genomes, and 8 mango genomes. Upon comparing the reference genomes with chromosomes assembled by various software combinations, we found that chromosomes assembled by Flye and Canu demonstrated higher completeness and accuracy compared to those assembled by wtdbg2. Both Hi-C sequence alignment software tools, HiC-Pro and HiCUP, showed similar performance in chromosome assembly outcomes, with no significant differences. In the context of chromosome-anchoring programs, both ALLHiC and LACHESIS proved effective in accurately clustering contigs to different groups. However, ALLHiC exhibited marked superiority in sorting and orienting the clustered contigs to reconstruct the correct chromosomes. These results not only underscore the importance of selecting appropriate assembly software to ensure high-quality genome reconstruction but also provides valuable insights for researchers in choosing the optimal tools for chromosome assembly, thereby enhancing the accuracy and efficiency of genomic research in various species.

In summary, the development of Chrom-pro has introduced a new tool in the field of genome assembly. Chrom-pro not only simplifies the chromosome assembly process but also incorporates multiple functions, such as genome quality assessment, comparative genomics analysis, and structural variation detection, providing comprehensive support for diverse genomic studies. Although extensive validation of Chrom-pro on large genomes such as the human genome remains untested, research utilizing similar software modules in chrom-pro has shown the capacity to handle large chromosomal genome assembly^70^, suggesting a wide range of potential applications in future genomic research.

## Author contributions

Conceptualization, W.S., H.J., S.L, and Y.H.; the development of Chrom-pro program and data analysis, W.S., T.Y., and S.L.; writing-draft preparation, W.S. and H.J.; writing-review and editing, H.J., W.S., Y.L., W.P., Y.H., Y.D., Y.Y., D.S., and HL. J.; supervision, H.J., W.P. and Y.H.. All authors have read and agreed to the published version of the manuscript.

## Supplementary Materials

**Figure S1.**
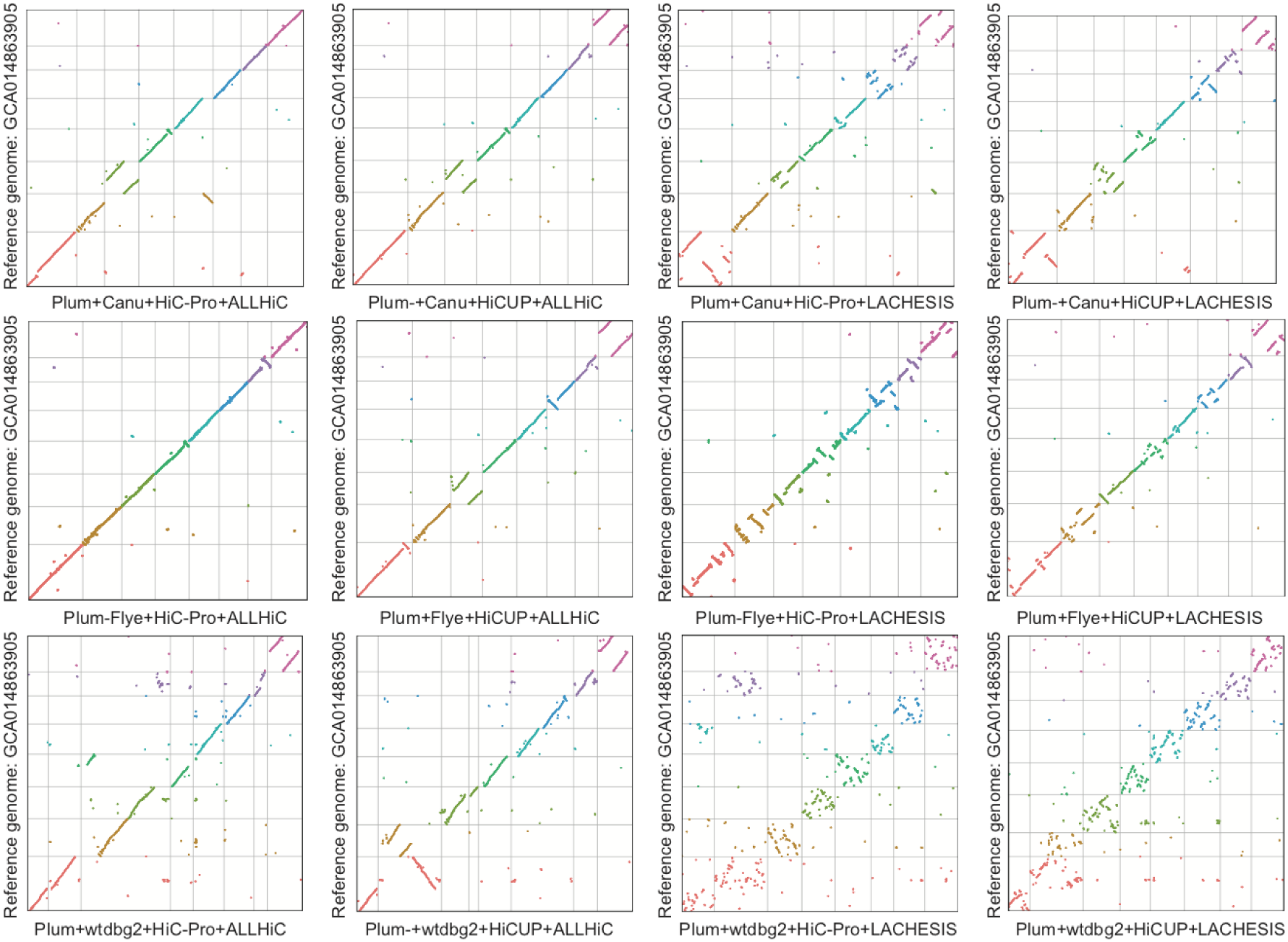
Collinearity analysis between the reference genome and plum chromosomes assembled by distinct software combinations.

**Table S1.** Sequencing data and the assembly methods of the reference genomes for Pufferfish, Mango, and Plum.

|  | Plum | Pufferfish | Mango |
| --- | --- | --- | --- |
| NGS | 26. 6Gb (85X) | 53. 43Gb (136X) | 7. 6Gb (21X) |
| TGS | 53Gb (170X) | 28. 97Gb (74X) | 86. 5Gb (240X) |
| Hi-C | 59. 1Gb (190X) | 46. 39Gb (118X) | 33. 8Gb (90X) |
| TGS assembler | FALCON | FALCON-unzip | Canu |
| Error correction | Quiver and Pilon | Arrow | Pilon |
| Haplotigs removing | Purge Haplotigs | Purge Haplotigs | Purge Haplotigs |
| Hi-C mapping | HiCUP | BWA | BWA |
| Chromosome anchoring | LACHESIS and Juicebox | BioNano scaffolding and SALSA2 | Juicer and 3d-dna and ALLMAPS |
| BUSCO complete | 96. 30% | 96. 60% | 98. 20% |

**Table S2.** Software used in chromosome assembly with Chrom-Pro for Pufferfish, Mango, and Plum.

|  | Plum | Pufferfish | Mango |
| --- | --- | --- | --- |
| TGS assembler | Flye | Canu | Canu |
| Error correction | Racon and Pilon | Racon and Pilon | Racon and Pilon |
| Haplotigs removing | Purge_dups | Purge_dups | Purge_dups |
| HiC mapping | HiC-Pro | HiC-Pro | HiC-Pro |
| chromsome anchor | ALLHiC | ALLHiC | ALLHiC |

